# Brassinosteroids Promote Parenchyma Cell and Secondary Xylem Development in Sugar Beet (*Beta vulgaris L.*) Root

**DOI:** 10.1101/2020.08.18.255976

**Authors:** Wei Wang, Yaqing Sun, Guolong Li, Shaoying Zhang

## Abstract

Increasing crop yield has always been an important goal in agriculture. Brassinosteroids (BRs) are growth-promoting steroid hormones with vital roles in many root developmental processes. Sugar beet (*Beta vulgaris L.*) is a root crop with a tertiary root structure. The differentiation of vascular bundles and division of cambial cells increases root diameter. However, little is known about how BRs regulate the transverse growth of beetroot. Therefore, sugar beet with eight leaves was grown in medium containing epibrassinolide or brassinazole, an inhibitor of brassinosteroid biosynthesis. BRs increased the spacing between the cambial rings by increasing the size of parenchyma cells between the rings and ultimately increasing root diameter. BRs also promoted secondary xylem differentiation. These findings indicate that brassinosteroids function in transverse development in beetroot.

## Introduction

Brassinosteroids (BRs) are ubiquitous plant hormones that have been used widely in agriculture since the 1970s (Divi and Krishna, 2009). Recent analyses have revealed that BRs are involved in many aspects of root development, including cell elongation, maintenance of meristem size, root hair formation, lateral root initiation, and the gravitropic response (Wei and Li, 2016). In Arabidopsis, a low BR concentration stimulated root elongation in wild-type plants by up to 50% and by up to 150% in BR-deficient mutants, such as *dwf*1-6. The root growth-promoting effect of BRs appears to be largely independent of auxin and gibberellin (Müssig et al., 2003). In roots, cell proliferation and post-mitotic cell enlargement form a developmental gradient along the apical–basal axis that eventually determines their length (Petricka et al., 2012). Moreover, cell enlargement is regulated by the extensibility of the surrounding wall (Cosgrove, 2005; Fujita et al., 2011; Wolf et al., 2012). Interestingly, recent chromatin-immunoprecipitation microarray (CHIP-chip) experiments identified several BR-regulated BRASSINAZOLE-RESISTANT1(BZR1) target genes that play vital roles in cell wall biosynthesis, such as cellulose synthase 6 (CESA6), xyloglucan endotransglycosylase/hydrolases (XTH), and pectinesterases, which ultimately promote cell elongation (Sun et al., 2010).

During the first year of the biennial life cycle of the sugar beet, root swelling and sucrose accumulation are the ultimate goals (Elliott and Weston, 1993). The beetroot is a crucial organ in sugar beet and accounts for 30% of the global sucrose output (Zhang *et al*., 2017). When a seedling becomes established, the plant enters a period of leaf initiation with very little root growth. At the 8–10 leaf stage, the leaves and root grow simultaneously; eventually, roots comprise the major proportion of the total plant dry weight (Elliott and Weston, 1993; Bellin *et al*., 2007). The increase in girth of the tap root results from the activity of the cambia. The innermost cambium is produced between the primary xylem and phloem. Subsequent cambia are initiated centrifugally in the outer portion of the previous ring (Milford, 1973). Although 12–15 cambial rings can form, the greatest contributions to root development are from rings 1 and 2, while rings 3–8 show progressively less activity. Rings 1–6 make up approximately 75% of the storage root (Elliott and Weston, 1993). Therefore, the rings that contribute most to the final yield of the root were already present when the plants had produced 12–13 leaves. The ringed structure of the beetroot in transverse section results from the development of parenchymatous zones and vascular bundles that contain xylem toward the inside and phloem toward the outside. However, few studies have examined whether BR promotes transverse growth by alternating the size or number of parenchyma cells in sugar beet.

In this study, we treated sugar beet with BR and brassinazole (Brz) to clarify the role of BR in the transverse growth of beetroot. This study offers a theoretical basis for further work on the molecular mechanisms of beetroot growth.

## Results

### BR increased the root diameter

Figure 1a shows sugar beets treated with BR, Brz, or water (control group) for 10 days. Compared with the controls, BR-treated plants had a significant (*P* < 0.01) increase in root diameter, while Brz-treated plants were not significantly altered (Fig. 1b). Therefore, BR increased the root diameter.

**Figure 1.**
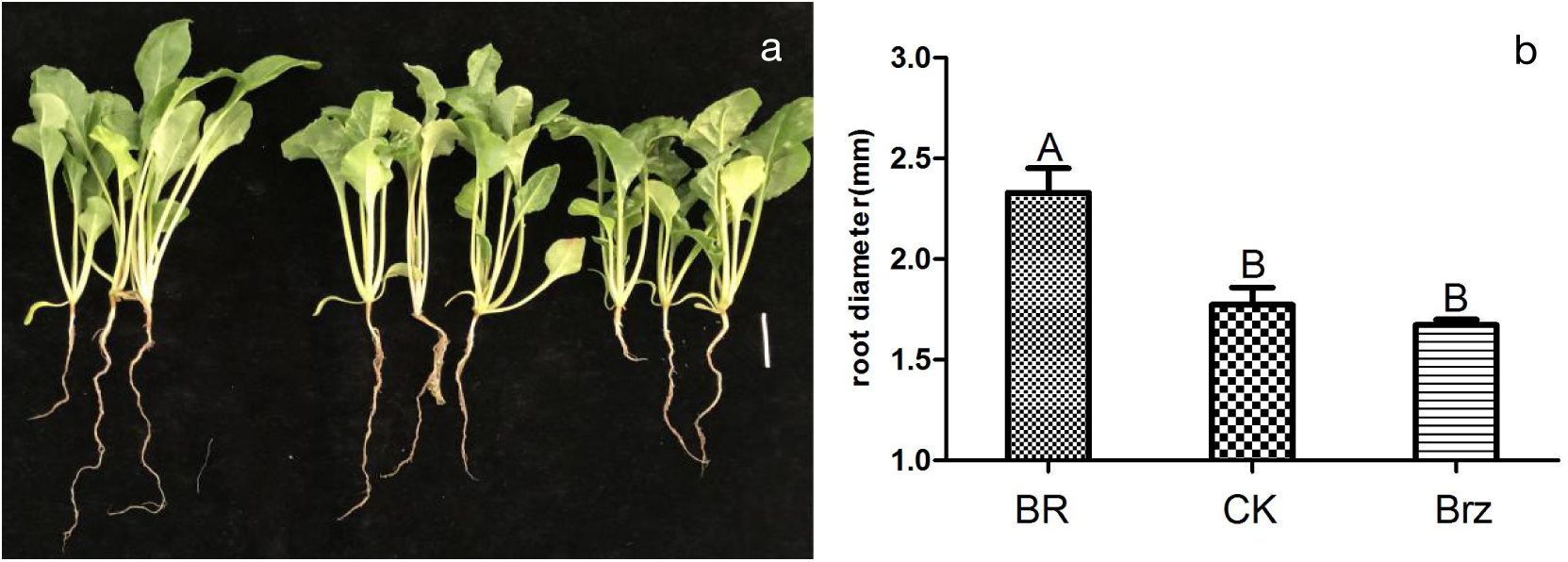
The effects of exogenous BR and Brz on sugar beet. (a) shows the morphological phenotype of sugar beet treated with BR,water or Brz. From left to right, BR(0.1mg/L)-treated,control and Brz(20µmol/L)-treated plants are shown. Sugar beet were grown for 10d. Scale bar equal to 3cm. (b) The figure illustrates the root diameter treated with BR,water or Brz in 10d. A B represent p<0.01.

### BR increased the spacing between cambial rings

To assess how root structure influences root diameter, we examined cross-sections of root apex treated with BR, Brz, or water (Figs. 3 and 4). After 10 days, all the experimental groups had four cambial rings. The second and third cambial rings, which contain the most vigorous cambial cells, were examined to assess the role of BR. Compared with the controls, BR-treated plants had a significant (*P* < 0.05) 37.5% increase in spacing between the first and second cambial rings (Fig. 2). Compared with the Brz-treated group, BR-treated plants had an increase of 54.5%. The spacings between the second and third rings in BR-treated plants were increased (*P* < 0.01) by 8% and 22.9%, respectively, compared with the control and Brz-treated plants (Fig. 2). Therefore, BR increases the spacing between cambial rings.

**Figure 2.**
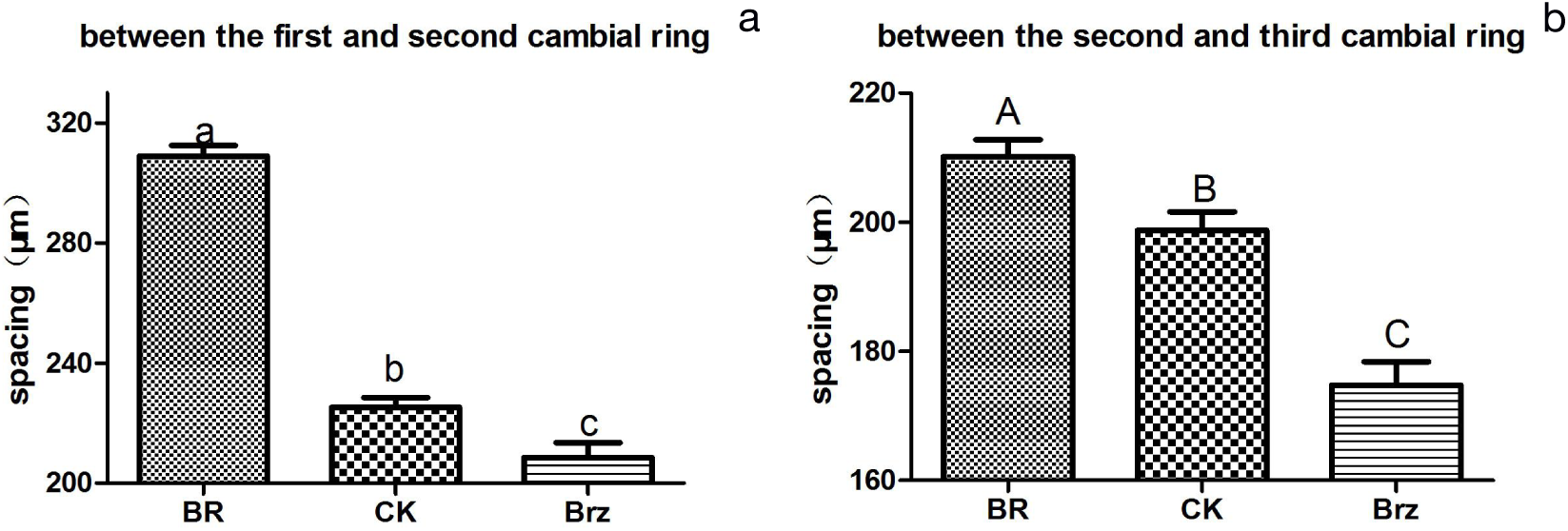
The effect of exogenous BR on spacing between cambial rings in beetroot. (a) The spacing between the first and second cambial rings in beetroot treated with BR, water or Brz. a b c represent p<0.05. (b) The spacing between the second and third cambial rings. A B C represent p<0.01.

**Figure 3.**
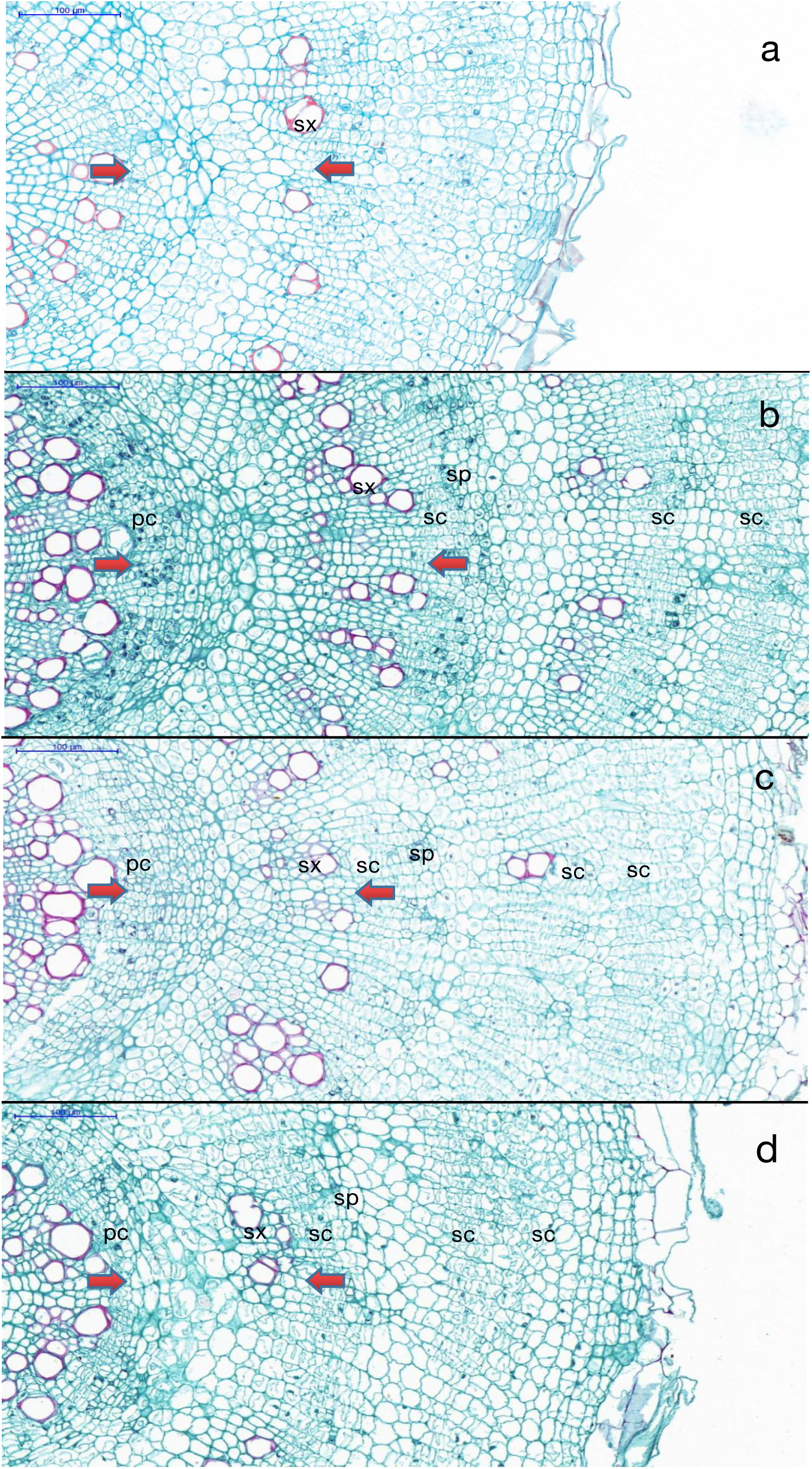
The effect of exogenous BR on the spacing between the first and second cambial rings in beetroot. Root cross-sections were stained with Safranin and Astra Blue. The distance between the two arrows indicates the spacing between cambial rings. pc,primary cambium;sx,secondary xylem;sc,secondary cambium;sp,secondary phloem. (a)root cross-section of control before the treatment.(b)root cross-section of BR(0.1mg/L)-treated sugar beet grown for 10d.(c)root cross-section of control sugar beet grown for 10d.(d)root cross-section of Brz(20µmol/L)-treated sugar beet grown for 10d. Bar=100µm.

**Figure 4.**
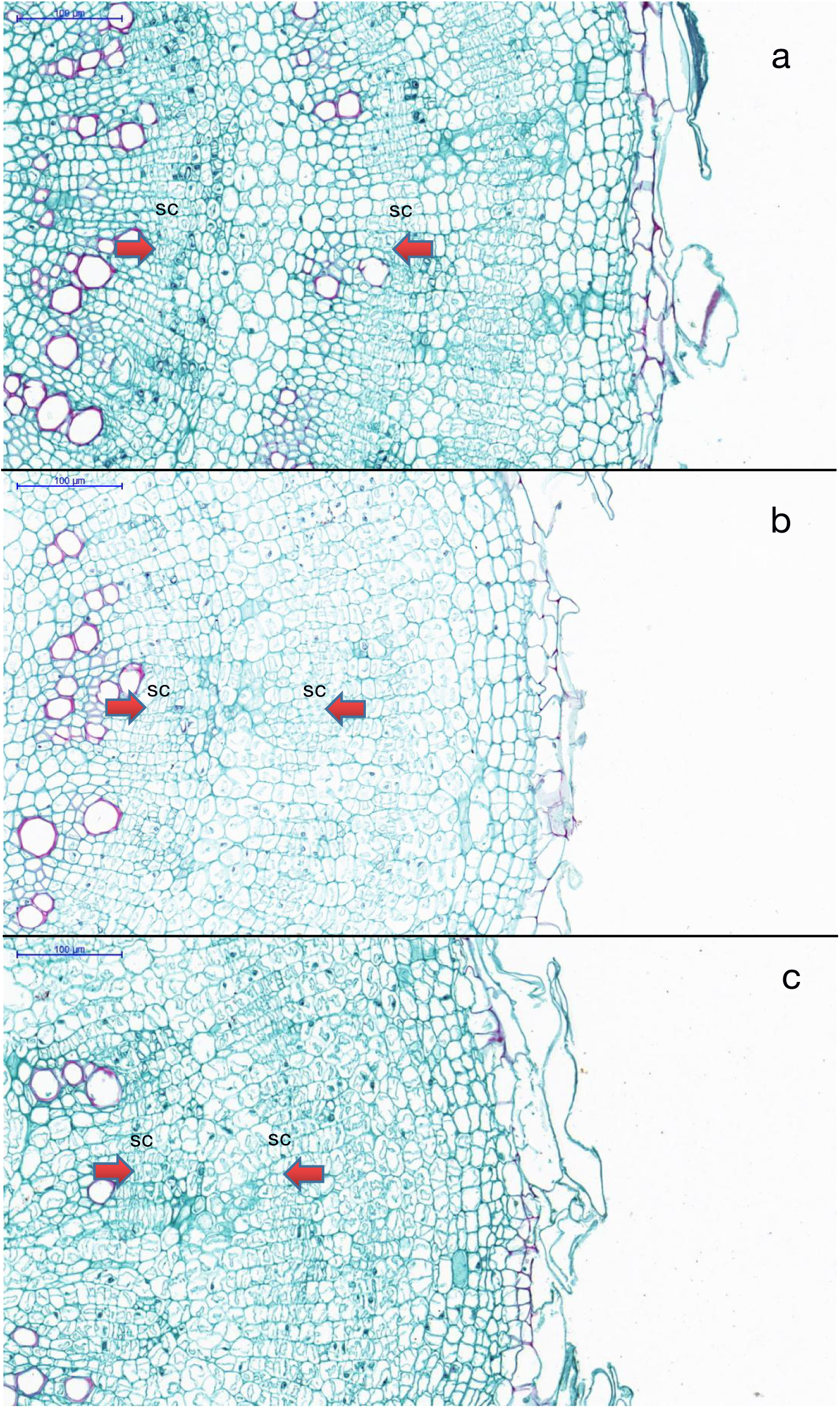
The effect of exogenous BR on the spacing between the second and third cambial rings in beetroot. Root cross-sections were stained with Safranin and Astra Blue. The distance between the two arrows indicates the spacing between cambial rings. sc,secondary cambium; (a)root cross-section of BR(0.1mg/L)-treated sugar beet grown for 10d.(b)root cross-section of control sugar beet grown for 10d.(c)root cross-section of Brz(20µmol/L)-treated sugar beet grown for 10d. Bar=100µm.

### BR increased the size of parenchyma cells

Figure 5 shows the parenchyma cells between the first and second cambial rings in plants treated with BR, Brz, or water. In BR-treated plants, the parenchyma cells between the first and second rings were significantly larger (23.8%). In comparison, parenchyma cells between the first and second rings in Brz-treated plants were 19.9% smaller (*P* > 0.05) (Fig. 5a–d). Compared with the controls, the layers of parenchyma cells between the first and second rings were significantly larger in BR-treated plants, but not Brz-treated plants (Fig. 5a–c, e).

**Figure 5.**
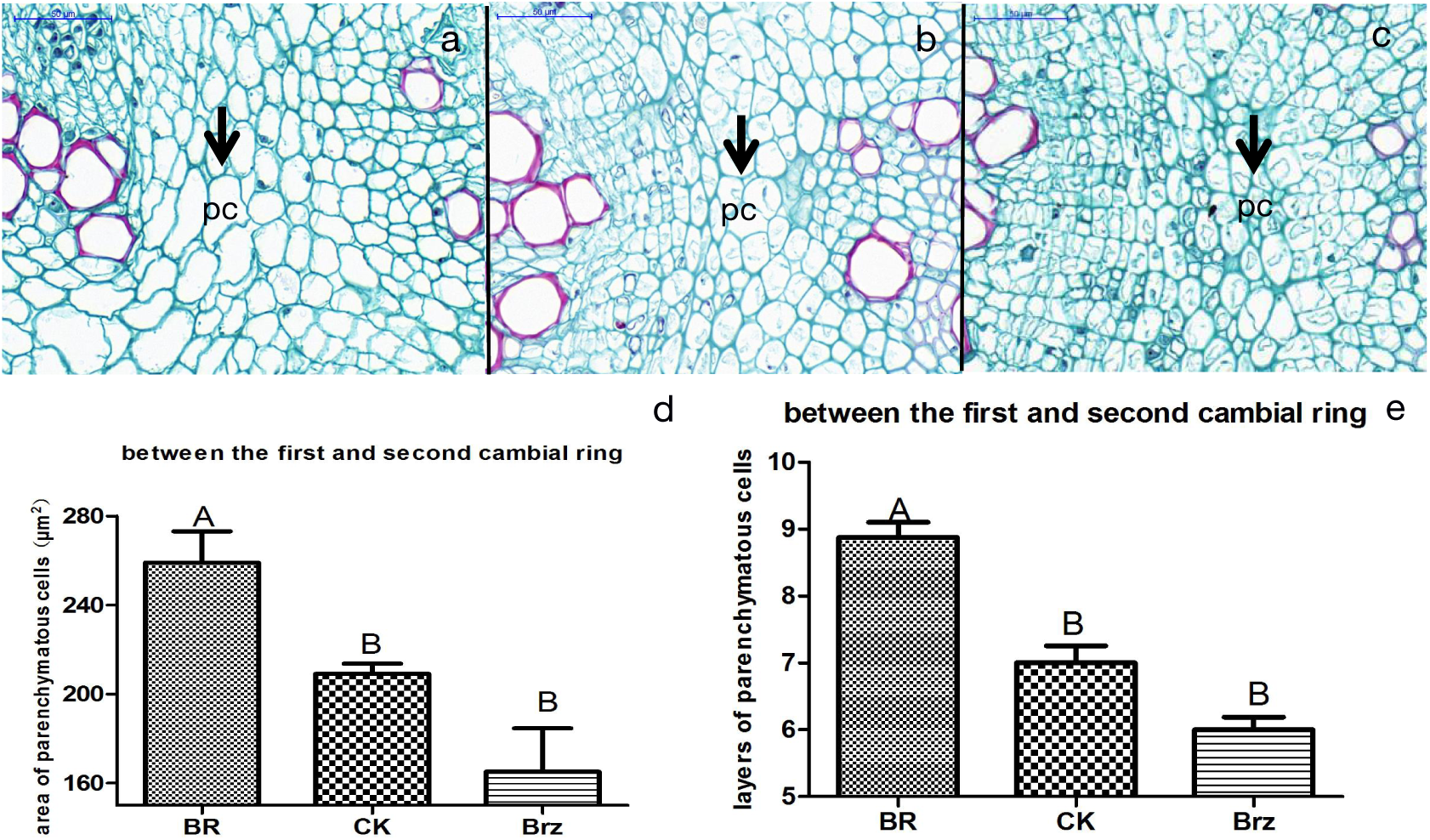
The effect of exogenous BR on parenchyma cells between the first and second cambial rings in beetroot. Root cross-sections were stained with Safranin and Astra Blue. The arrow represents the parenchyma cells. pc,parenchyma cell. (a)root cross-section of BR(0.1mg/L)-treated sugar beet grown for 10d.(b)root cross-section of control sugar beet grown for 10d.(c)root cross-section of Brz(20µmol/L)-treated sugar beet grown for 10d. (d)reveals the area of the parechyma cells between the first and second cambial rings after treatments for 10d. (e) illustrates the layers of the parechyma cells between the first and second cambial rings after treatments for 10d.A B represent p<0.01. Bar=50µm in (a,b,c).

While BR treatment increased the size of parenchyma cells between the second and third cambial rings by 6.7% compared with the controls, Brz treatment decreased the size by 6.5% (both *P* < 0.01; Fig. 6a–d). The number of layers of parenchyma cells did not differ among the BR-treated, Brz-treated, and control groups (Fig. 6a–c, e). Accordingly, BR increases the size of parenchyma cells.

**Figure 6.**
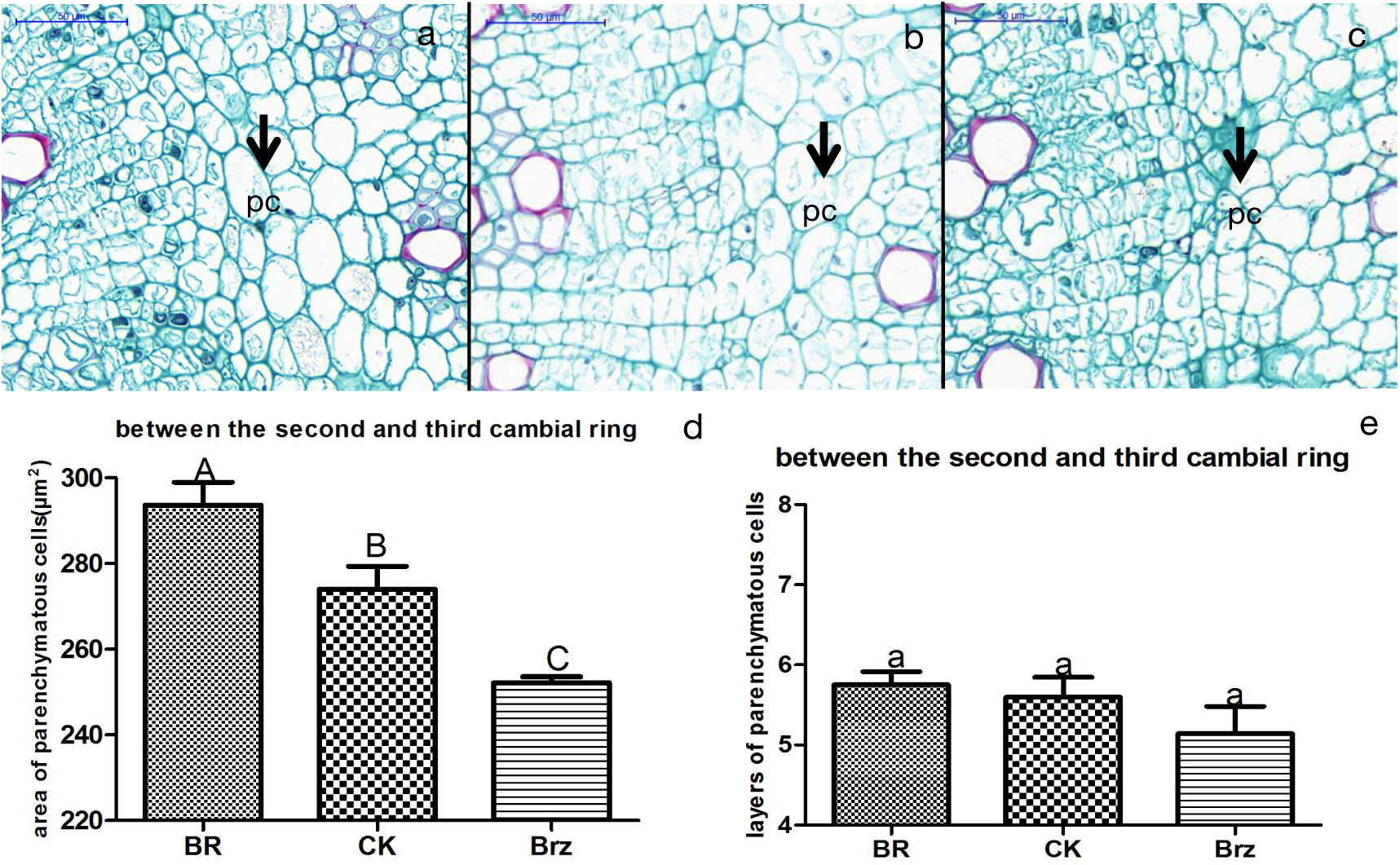
The effect of exogenous BR on parenchyma cells between the second and third cambial rings in beetroot. Root cross-sections were stained with Safranin and Astra Blue. The arrow represents the parenchyma cells. pc,parenchyma cell. (a)root cross-section of BR(0.1mg/L)-treated sugar beet grown for 10d.(b)root cross-section of control sugar beet grown for 10d.(c)root cross-section of Brz(20µmol/L)-treated sugar beet grown for 10d. (d)reveals the area of the parenchyma cells between the second and third cambial rings after treatments for 10d. (e) illustrates the layers of the parenchyma cells between the second and third cambial rings after treatments for 10d. A B C represent p<0.01. a represent p<0.05.Bar=50µm in (a,b,c).

**Figure 7.**
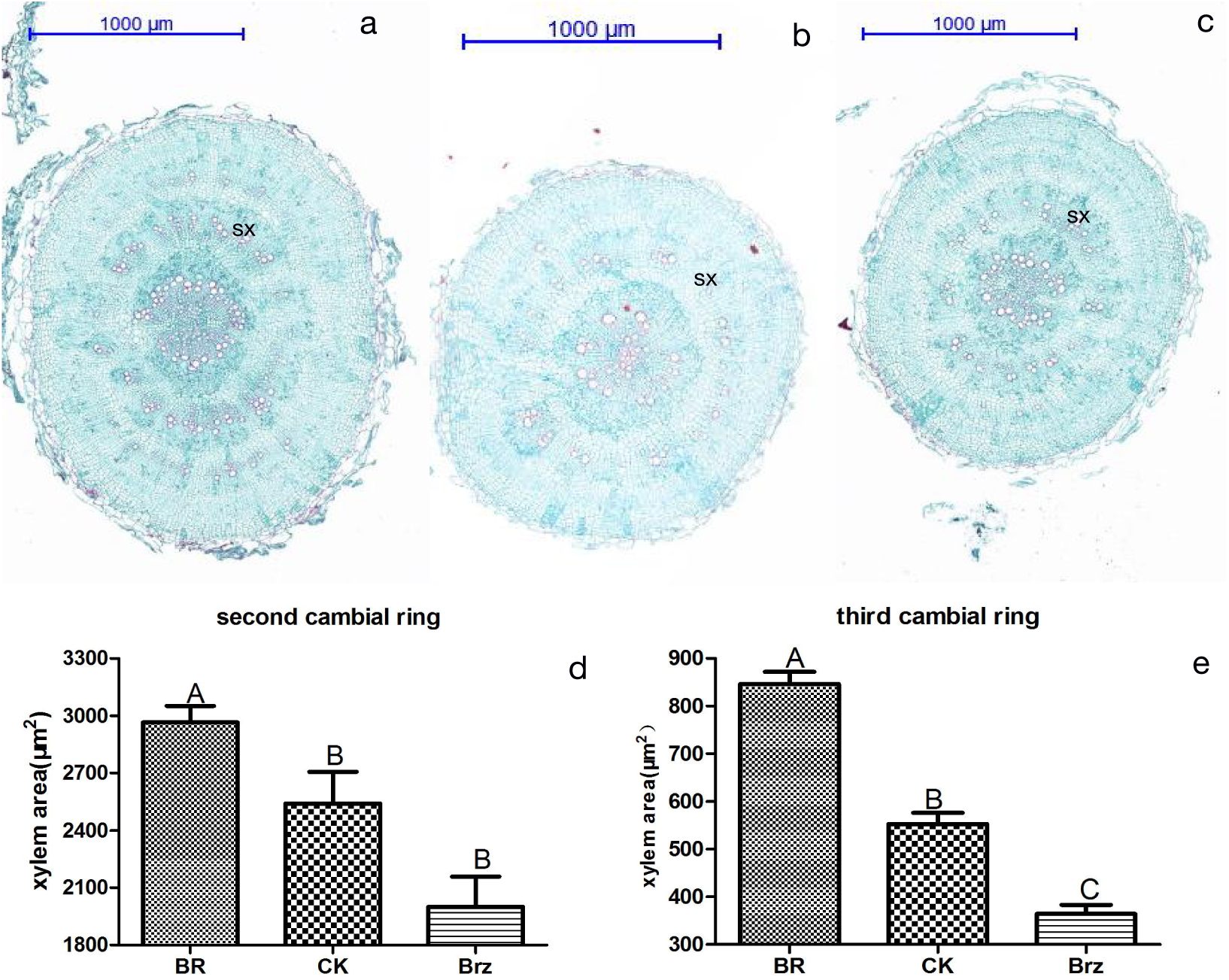
The effect of exogenous BR on secondary xylem in beetroot. Root cross-sections were stained with Safranin and Astra Blue. sx, secondary xylem.(a)root cross-section of BR(0.1mg/L)-treated sugar beet grown for 10d.(b)root cross-section of control sugar beet grown for 10d.(c)root cross-section of Brz(20µmol/L)-treated sugar beet grown for 10d. (d) reflects the area of the secondary xylem at the second cambial ring after treatments for 10d. (e) illustrates the area of the secondary xylem at the third cambial ring after treatments for 10d. A B C represent p<0.01. Bar=1000µm in (a,b,c).

### BR promoted the development of secondary xylem

The areas of secondary xylem in the second and third cambial rings were calculated using CaseViewer. In BR-treated plants, these areas increased significantly (15.58% and 53.15%, respectively; *P* < 0.01) compared with the controls. Hence, BR promoted development of the secondary xylem.

## Discussion

There have been many recent reports on how BRs influence plant root development. At certain concentrations, BRs decrease the root length by inhibiting apical meristem growth. For example, in wheat, 0.1 and 1 nmol/L 24-eBL promoted root growth, while 10 nmol/L 24-eBL inhibited root growth. However, the research on the effects of BRs on plant roots has focused on longitudinal growth, while transverse growth is rarely studied. In sugar beet, studying root swelling is important. Comparing the three treatments, BR promoted root swelling by increasing the spacing between the cambial rings, which depends on the number and size of parenchyma cells between the rings. BR produced larger parenchyma cells, but had no significant effect on the number of parenchyma cells, which is consistent with the findings that BR mediates root cell elongation (Wei and Li, 2016).

Vascular bundles are necessary for the growth and development of higher plants, as they transport plant materials. BR affects xylem development. When some BR synthesis genes were mutated, the xylem decreased and phloem increased in vascular tissue (Choe, 1999). Brz treatment inhibited the secondary xylem in cress (Nagata, 2001). Jiaxing (2012) found that the *TCP1* gene, which regulates BR synthesis, is involved in the differentiation and formation of vascular epigenetic xylem in Arabidopsis. We found that the area of secondary xylem in beetroot increased with BR treatment, but decreased with Brz treatment. The area of xylem in the third cambial ring of beetroot treated with BR increased by 53.15% compared with the controls. Therefore, BR promotes beetroot xylem.

## Conclusion

On treating sugar beet at the eight-leaf stage with BR, Brz, or water, BR increased the beetroot diameter. Histological analyses revealed that BR increases the spacing between the cambial rings by increasing the size of parenchyma cells between the cambial rings. BR also promotes the differentiation of xylem.

## Material and Methods

### Plant materials and growth conditions

The BS02 sugar beet cultivar used in this study was bred in our laboratory. Seeds were cultivated in the Inner Mongolia Agricultural University phytotron (Hohhot, Inner Mongolia, China) and sown in vermiculite with one seedling per 8 × 8 × 8 cm3 float tray. Hoagland solution was added every 14 days and the plants were grown at 22℃ and under 16 h light/8 h dark conditions.

### BR and Brz treatments

The eight-leaf plants were treated with 0.1 mg/L epibrassinolide (epiBL; Sigma, USA) or 20 µmol/L Brz (Sigma,USA). The epiBL and Brz solutions were prepared by dissolving the solute in DMSO and diluting with distilled water. The control group was treated with water. On day 10, the morphological and physiological parameters of all plants were measured and the root tissues were harvested for histological analyses. Roots were collected from three biological replicates.

### Histological parameters

Samples were collected from the root 1 cm from the root apex. All specimens were fixed in 70% formaldehyde acetic acid for 24 h and dehydrated in an increasing ethanol series. Transverse 12-µm-thick sections were obtained with a rotating microtome, stained with Safranin and Astra Blue, and mounted on paraffin (Maia et al., 2018). The slides were observed under a microscope (Pannoramic DESK, Hungary) and photomicrographed. The images were analyzed with CaseViewer. The following histological parameters were evaluated: size of the xylem and parenchyma cells and spacing between the cambial rings.

### Data analysis

The differences between the mean values were assessed at 5% and 1% probability levels using SPSS (ver. 18.0). The figures were made with GraphPad Prism (ver. 8.0).

The English in this document has been checked by at least two professional editors, both native speakers of English. For a certificate, please see: http://www.textcheck.com/certificate/jjxQVQ

